# A comparative transcriptional landscape of two castor cultivars obtained by single-molecule sequencing comparative analysis

**DOI:** 10.1101/2020.09.30.320101

**Authors:** Wei Zhou, Yaxing Zhou, Guoli Zhu, Yun Wang, Zhibiao He, Zhensheng Shi

## Abstract

**Background and Objectives:** Castor (*Ricinus communis* L.) is an important non-edible oilseed crop. Lm type female strains and normal amphiprotic strains are important castor cultivars, and are mainly different in inflorescence structures and leaf shapes. To better understand the mechanisums underling these differences at the molecular level, we performed comparative transcriptional analysis.

**Materials and Methods:** Full-length transcriptome sequencing and short-read RNA sequencing were employed.

**Results:** A total of 76,068 and 44,223 non-redundant transcripts were obtained from high-quality transcripts of Lm type female strains and normal amphiprotic strains, respectively. In Lm female strain and normal amphiprotic strains 51,613 and 20,152 alternative splicing events were found, respectively. There were 13,239 transcription factors identified from the full-length transcriptomes. Comparative analysis showed great different gene expression of common and unique transcription factors between the two cultivars. Meanwhile, functional analysis of isoform was conducted. Full-length sequences were used as a reference genome, and short-read RNA sequencing analysis was performed to conduct differential gene analysis. Furthermore, the function of DEGs were performed to annotation analysis.

**Conclusions:** The results revealed considerable difference and expression diversity between two cultivars, well beyond what was reported in previous studies, likely reflecting the differences in architecture between these two cultivars.

**Highlight:** Using the full-length transcriptome sequencing technology, we performed comparative analysis of transcription factors of two castor cultivars, analyzed alternative splicing events, and identified their lncRNAs.

## Introduction

The castor plant (*Ricinus communis* L.), originated in Africa, is an annual or perennial dicotyledonous. High ricinoleic acid content (80-90%) and high fatty acid content (more than 45%) in seed oil make oit one of the most important non-edible oilseed crops, and has attracted much attention of chemists, biologists and medical scientists [1]. The inflorescence of common castor plants is gradient monoecious raceme, with staminate flowers on the lower portion and female flowers at the apex[2]. Pistillate (bearing only female flowers) variations are bred to improve the seed yield. The Lm type castor is such a variety obtained by exposing castor seeds to ^60^Coγ.

With the development of sequencing technology, next-generation sequencing (NGS) has become an essential method for studies of genomes, epigenomes, and transcriptomes [1]. The NGS method has been used in many model and non-model plant species, and large-scale genome sequences and transcriptome data have been produced for deep analysis [1]. However, the deficiency of NGS, for example short reads, result in incompletely assembled transcripts limiting to better understand the transcriptomic data [1]. PacBio platform, based on the single-molecule real-time (SMRT) sequencing technology and providing longer and full-length transcripts without assembly, can provide better information to understand the full-length transcriptome, such as alternative splicing, fusion transcripts, alternative polyadenylation, novel genes and non-coding RNAs [1].

To gain an insight how sex is differentially regulated at the molecular level, in the present study, full-length transcriptomes of Lm type and normal castor cultivars were analyzed, and short-read RNA sequencing and single-molecule long-read sequencing were utilized to identify the differentially expressed genes and alternative splicing events between Lm type female strains and normal amphiprotic strains. Furthermore, the study will provide valuable data for future studies of sex determination on the castor plants.

## Materials and methods

Two cultivars, Lm type female plants with willow-shaped leaves and normal amphiprotic plants (Fig. 1), were grown at the Experimental Base of the Agricultural College of Inner Mongolia University for Nationalities, Tongliao City, Inner Mongolia Autonomous Region. The geographical position is between 42°15′-45°41′ north latitude and 119°15′-123°43′ east longitude. Ten plants of each cultivar were selected when the functional leaves grew to 1-6 cm (Lm type) or 2-15 cm (normal type) on August 2nd, 2018. The Lm type female plants and normal amphiprotic plants were designated as F01 and F02, respectively. For each cultivar around 10g of leaves and flowers were collected and frozen in liquid nitrogen and then stored at −80 °C for subsequent RNA isolation. For Illumina HiSeq X Ten platform, RNAs were extracted from 5g of frozen leaves or flowers with two repeats, and RNA-Seq library construction was performed following the instructions[3].

**Figure 1.**
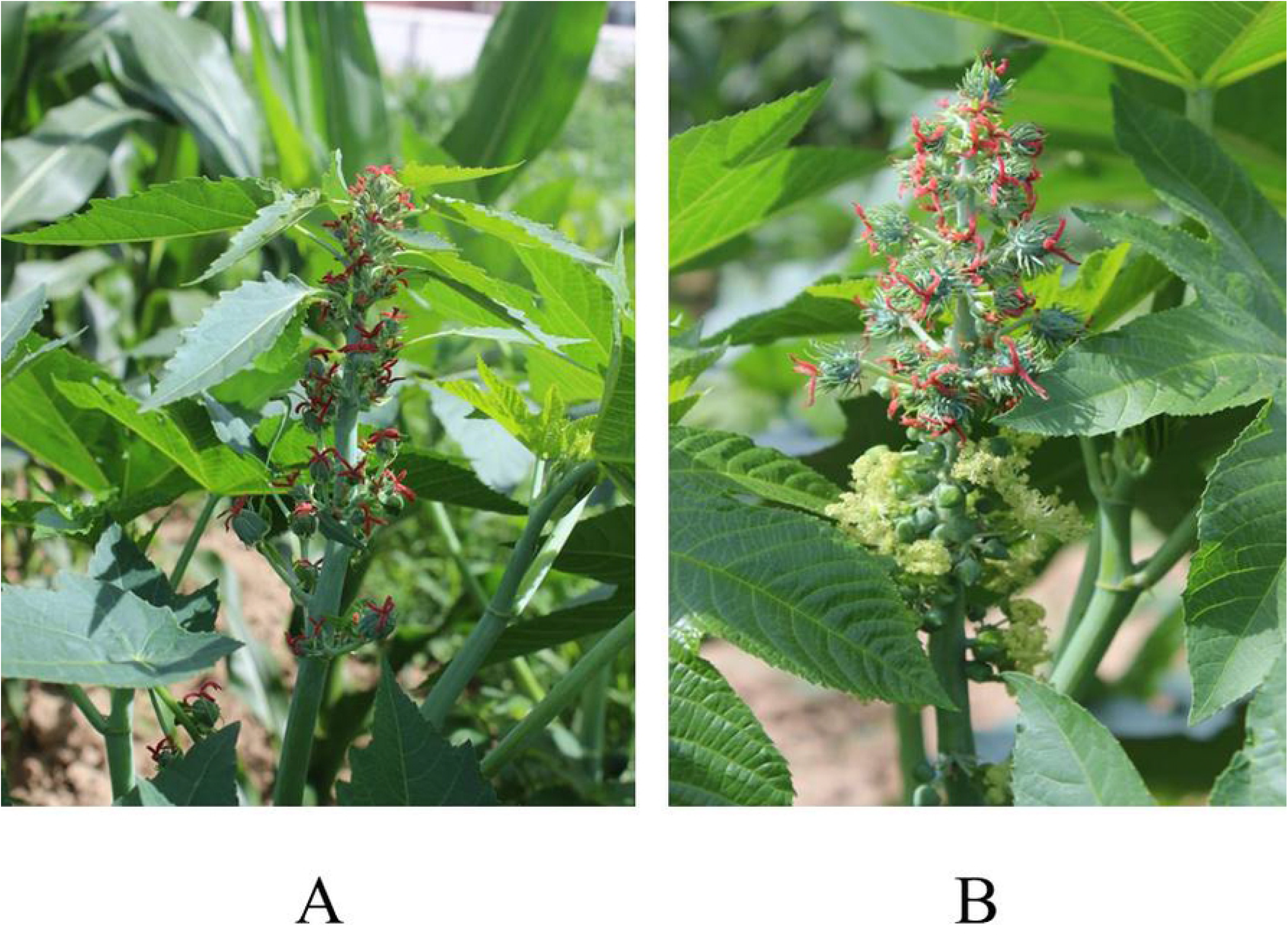
The castor plant R. communis L. (A) Lm type female strains with willow-shaped leaves. (B) Normal amphiprotic strains.

**Figure 2.**
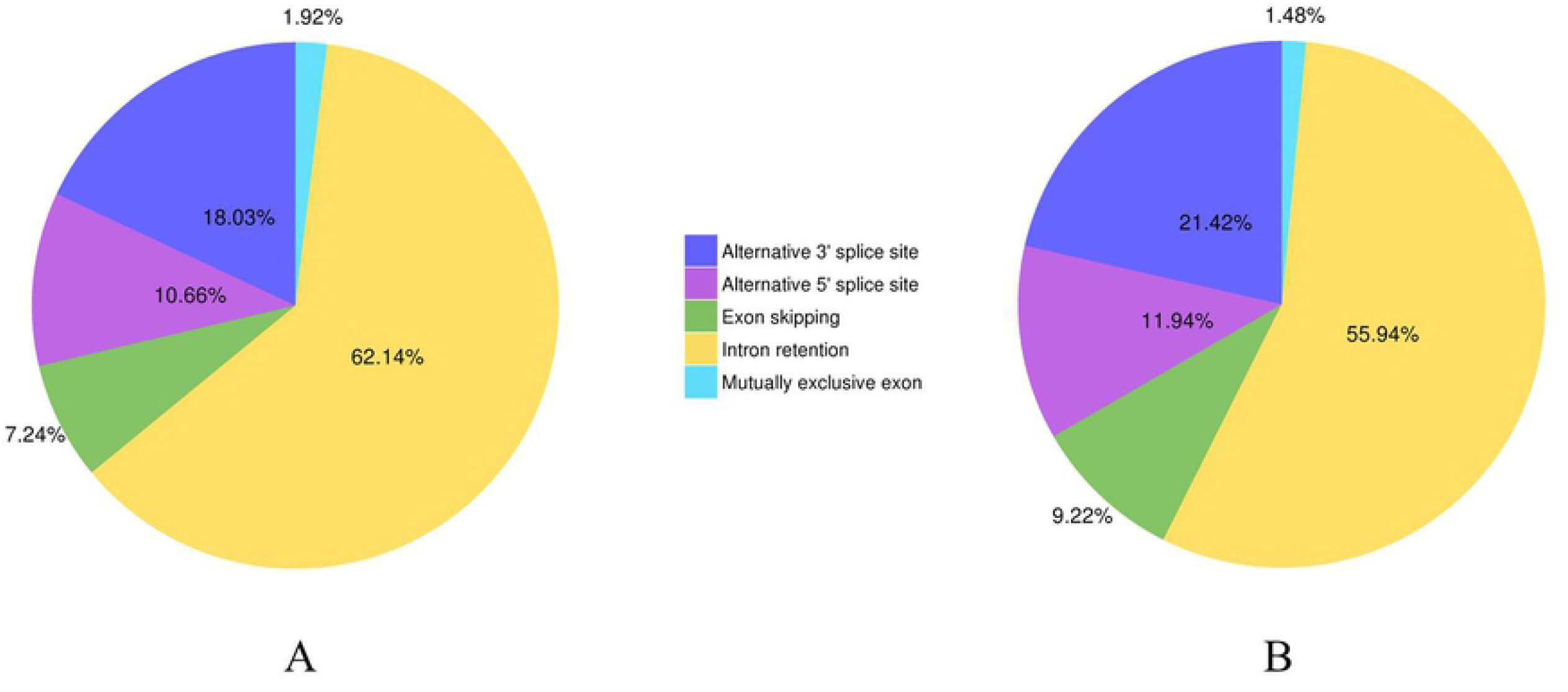
The statistics of different AS events of two cultivars. (A) AS events of Lm female strain. (B) AS events of normal castor.

### PacBio Library Construction and Sequencing

Total RNAs were extracted from 5g mixtures of leaves and flowers (two repeats). Poly(T) oligo-attached magnetic beads (Dynal) was used to purify mRNAs from about 3 µg total RNA. According to the protocol of PacBio RS II platform, cDNA was synthesized using the SMART PCR cDNA Synthesis Kit (Clontech, CA, USA), and then fractionized with BluePippin^®^ (Sage Science, Beverly, MA, USA). Then the final libraries were constructed using the Pacific Biosciences DNA Template Prep Kit (version 2.0). SMRT sequencing was performed on the Pacific Biosciences’ real-time sequencer using C2 sequencing reagents.

### Preprocessing of SMRT reads

The subreads were filtered using the standard protocol of the SMRT Analysis software suite (http://www.pacificbiosciences.com), and the reads of insert (ROIs) were obtained. After examining the poly(A) signals and 5’ and 3’ adaptors, full-length (FL) and non-full-length (nFL) reads were identified.

Consensus sequences were obtained from high-quality isoform sequences. The final transcriptome isoform sequences were filtered by removing the redundant sequences with the CD-HIT package (http://weizhong-lab.ucsd.edu/cdhit_suite/cgi-bin/index.cgi?cmd=cd-hit) to cluster and compare protein or nucleotide sequences.

### Alternative splicing analysis of Transcriptomes

To identify alternative splicing (AS) events, SpliceGrapher [4] were used to analyze the transcriptome-wide AS events. AS events were predicted from nonredundant transcripts. The prediction criterion is as following: the sequence should be greater than 1000 bp, AS gap should be greater than 100 bp, and at least 100 bp from the 3’-/5’-end, and there is a 5-bp overlap in the spliced transcript. Compared with the reference castor genome (http://castorbean.jcvi.org/), the full-length transcripts can be classified as derivation from the known genes and novel genetic loci.

Candidate coding regions were identified by TransDecoder (Broad Institute, Cambridge, MA, USA) from the final transcriptome isoform sequence. Sequences were searched using BLASTX [5] against the NCBI non-redundant protein and the UniProt with E-value cutoff at 1×10^−6^. To further distinguish protein-coding and non-coding RNAs, dbHT-Trans tool (v1.0)[6] were used for all PacBio transcripts.

Gene Ontology (GO) enrichments were analysed using the GOseq [7]. KEGG (http://www.genome.jp/kegg/) pathway analysis was implemented as reported [8].

### Short-read RNA sequencing analysis and quantification of gene expression

Clean reads were screened from raw sequencing reads by removing low-quality reads or reads containing adaptors or ploy-Ns. Sequences of clean reads were aligned to the full-length sequences.Differential expression analysis was performed with EBSeq package [9], with FDR < 0.05 and |log2 (fold-change) | ≥1.

## Results

### PacBio Iso-Seq Sequencing

SMRT sequencing generated 456,994 polymerase reads in total, and 26.25 Gb and 16.38 Gb clean reads were obtained from Lm female and normal castor cultivars, respectively. Under the conditions of full passes of ≥0 and quality of >0.80, 647,205 and 328,497 ROIs were obtained from two cultivars, respectively (Supplemental Table 1). In addition, 448,217 and 258,645 full-length non-chimeric sequences were identified from Lm and normal castors, respectively (Supplemental Table 2).

**Table 1.** The statistic results of ICE clustering.

| Samples | Number of consensus<br>isoforms | Average consensus isoforms<br>read length | Number of polished<br>high-quality isoforms | Number of polished<br>low-quality isoforms | Percent of polished<br>high-quality isoforms (%) |
| --- | --- | --- | --- | --- | --- |
| F01 | 223,929 | 2,552 | 154,517 | 68,417 | 69.00% |
| F02 | 138,066 | 2,056 | 105,536 | 32,086 | 76.44% |

**Table 2.** Statistics of annotations for genes of castor.

| Anno Database | Annotated Number |
| --- | --- |
| COG Annotation | 35,743 |
| GO Annotation | 61,390 |
| KEGG Annotation | 39,598 |
| KOG Annotation | 57,228 |
| Pfam Annotation | 63,089 |
| Swissprot Annotation | 62,336 |
| eggNOG Annotation | 83,826 |
| Nr Annotation | 85,286 |
| All Annotated | 85,322 |

The lengths of full-length cDNA in Lm female strain ranged from 281 to 11430 bp with an average length of 2702 bp. For normal castor train, full-length cDNAs showed an average length of 2192 bp, ranged from 303 bp to 9681 bp. The N50 values of those cDNAs were 3093 bp and 2408 bp in Lm and normal castor, respectively. Then, from 223,929 (Lm female cultivar) and 138,066 (normal castor) full-length consensuses cDNAs, 76,068 out of 154,517 (49%) and 44223 out of 105,536 (42%) high quality full-length consensuses were obtained, respectively. The ICE clustering results were shown in Table 1.

### Alternative splicing and polyadenylation

A total of 51,613 and 20,152 AS events were found in Lm and normal castors, respectively, including exon skipping (ES), intron retention (IR), alternative 3′ sites (Alt. 3′), alternative 5′ sites (Alt. 5′) and mutually exclusive exon. The results showed that intron retention (IR) was the foremost AS events, with 62.14% and 55.94% in two cultivars, respectively. The results of statistical analysis of different AS events of Lm female strain and normal castor were showed in Fig.2 and Supplemental Table 3.

### Comparative analysis of LncRNA and Transcription factors

Transcription factors which need to specifically bind to certain genes are essential for regulation of gene expression. A total of 13,239 encoded transcription factors were identified from the full-length transcriptome in two cultivars. Furthermore, we performed comparative analysis of common and unique transcription factors in the two cultivars. The main transcription factor types in Lm type castor include Rlk-Pelle-Dlsv, C3H, SNF2, MYB-related families. For normal castor, dominant transcription factors were Rlk-pelle-dlsv, camk-camkl-chk1 and MYB-related bHLH types. Although two cultivars shared some types of transcription factors, the expression of corresponding genes was completely different (Fig. 3). As the key regulators in biological processes, long non-coding RNA (LncRNA) is type of RNA that does not encode proteins [10]. A total of 858 lncRNAs were found in two cultivars using CPC, CNCI, CPAT and PFAM software [11]. Genomic distributions of LncRNAs were classified into 4 types, namely lincRNA, antisense-lncRNA, intronic-lncRNA and sense_lncRNA. The ratios of different types were waried greately, with 285 lincRNA, 58 antisense-lncRNA, 7 intronic-lncRNA and 166 sense_lncRNA in Lm type, and 60, 22, 3 and 49 in normal castor cultivar, respectively (Fig. 4).

**Figure 3.**
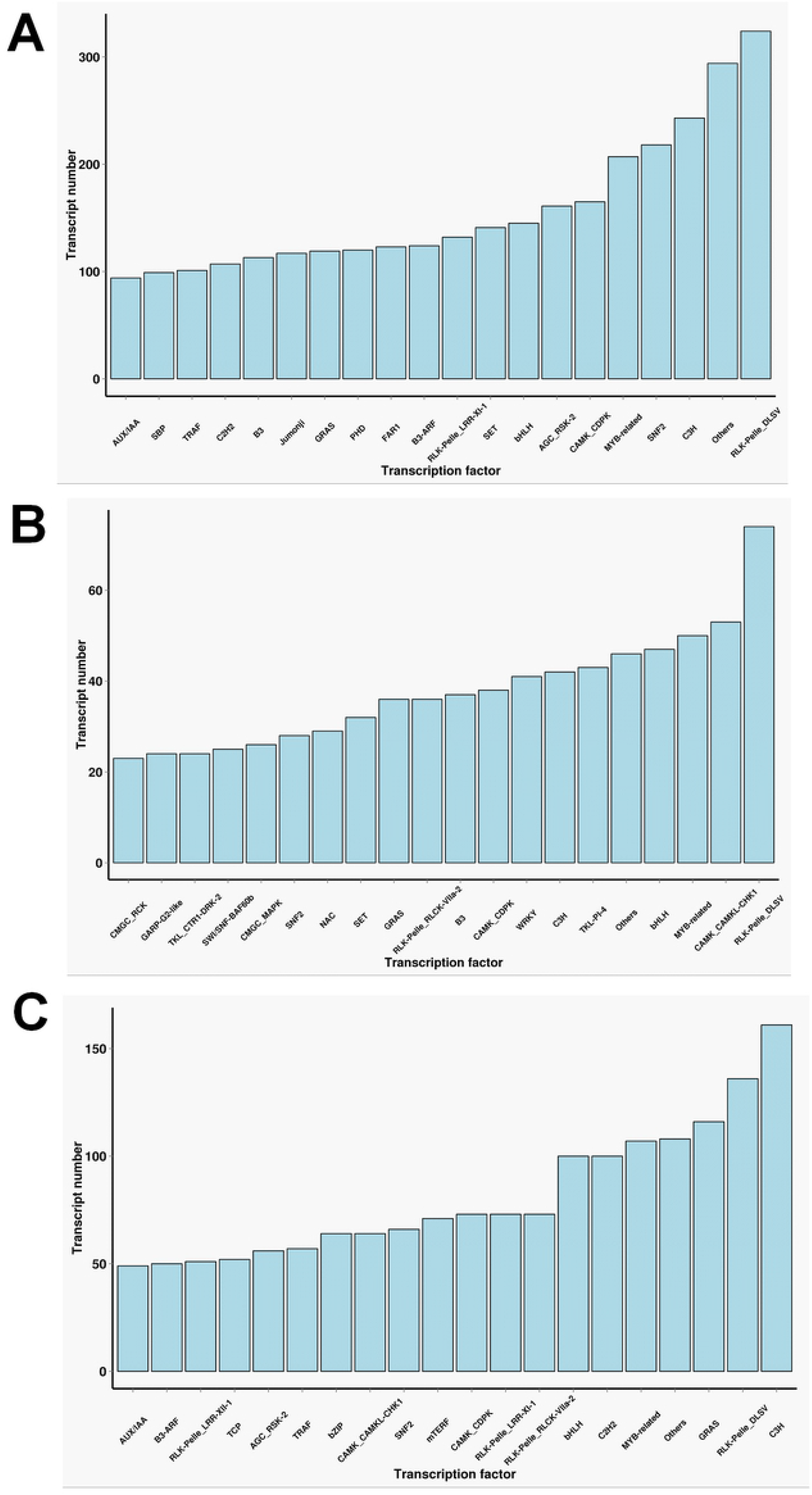
Statistics of transcription factors of two cultivars. **(A)** The number of transcription factor only in F01. **(B)** The number of transcription factor only in F01. **(C)** The number of transcription factor common in F01 and F02.

**Figure 4.**
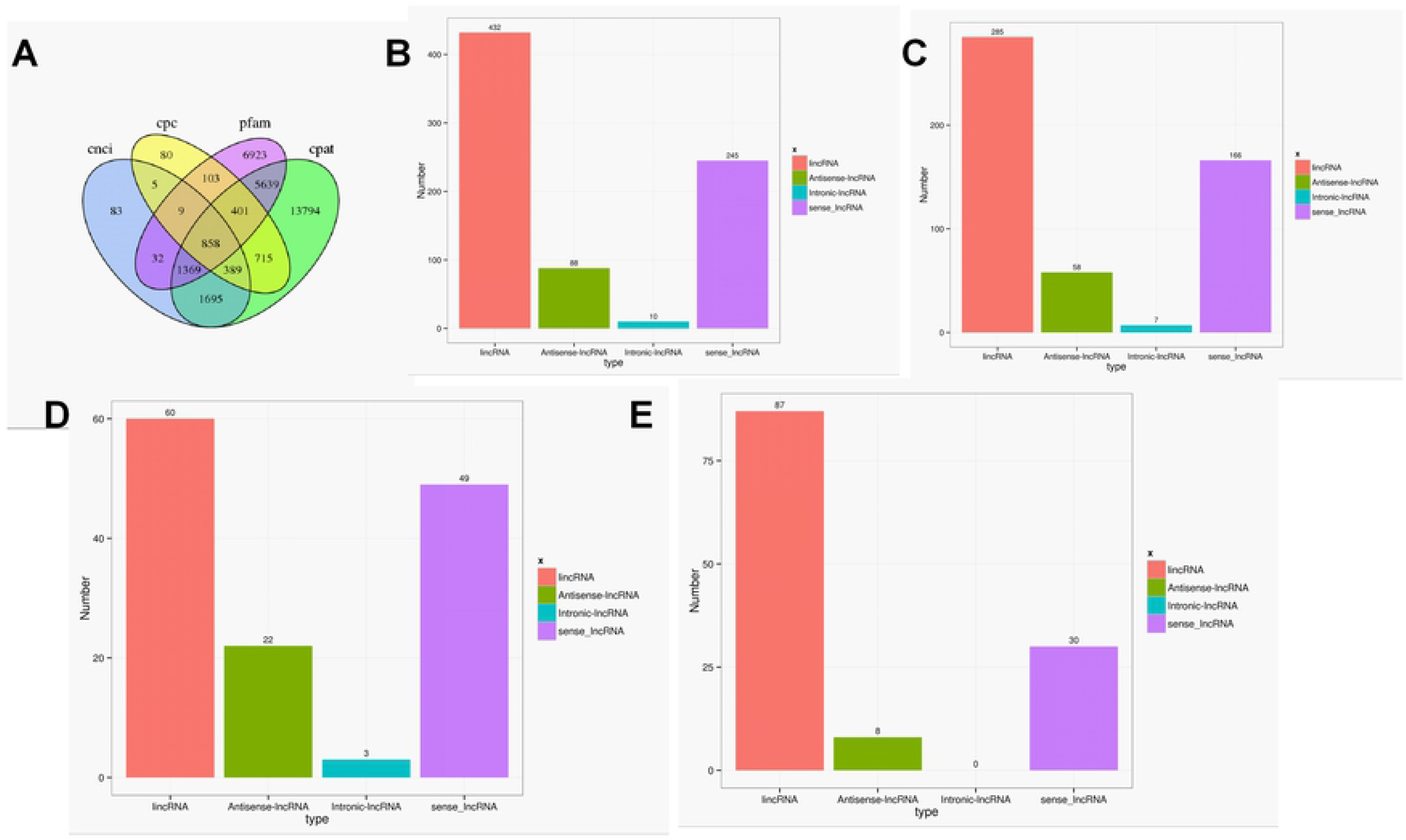
Statistics of LncRNA in two cultivars. (A) Venn diagram of LncRNAs using CPC, CNCI, CPAT and PFAM software. (B) Statistics of LncRNA types in genomic distributions. (C) Statistics of LncRNA types only in F01. (D) Statistics of LncRNA types only in F02. (E) Statistics of LncRNA types common in F01 and F02.

### Functional Annotation and Analysis of Isoform

For the functional annotation of gene isoforms, these genes were searched against Genbank NR, Swissprot, GO, COG, KOG, Pfam and KEGG database, and a total of 85,322 genes were annotated by seven databases. Among them, 85,286 genes (99.96 %) were aligned to the NR, and 62,336 genes were matched to the SWISS-PROT (Table 2). Approximately 79.21% of genes were aligned to *R. communis*, followed by *Jatropha curcas* (8.27%) (Fig. 5A).

**Figure 5.**
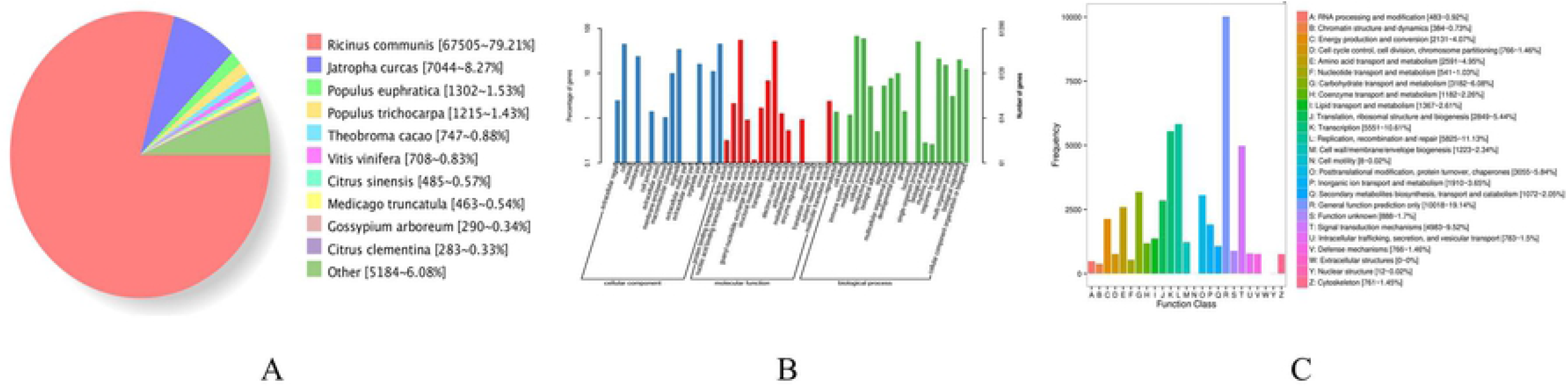
Statistics of the gene annotation in castor R. communis. (A) Nr homologous species distribution statistics. (B) GO annotation classification statistics. (C) COG annotation classification statistics.

GO annotation system is a directed acyclic graph, including three categories: biological process (BP), molecular function (MF), and cellular component (CC). In this study, GO analysis was conducted using Blast2GO, detailed GO distributions in GO categories were shown in Fig. 5B. The vast majority of genes were in cell or cell part in the cellular component. In the molecular function class, most of the genes were classified catalytic activity and binding. In the biological process class, genes of metabolic processes and cellular processes were the most. COG refers to Clusters of orthologous groups for eukaryotic complete genomes, and every protein in the database is assumed to be evolved from a common ancestor protein. In total 35,743 out of 85,286 genes were classified into 25 different COG categories (Fig. 5C), and genes with general functions were the largest category, followed by replication, recombination and repair, and transcription.

KEGG database is used to determine whether the genes were involved in specific metabolic or signal transduction pathways. In this study, a total of 125 KEGG pathways were identified. Several enriched pathways were involved in plant hormone signal transduction (ko04075), starch and sucrose metabolism (ko00500) and protein processing in the endoplasmic reticulum (ko04141) (Supplemental list 1).

### Functional comparative analysis of isoform

For further annotation analysis of gene function, the function of specific and common isoforms in two samples were analyzed systematically, indicating, although many of isoforms in the two samples were different, corresponding gene functions were similar. GO analysis showed that many of isforms enriched in the following items: metabolic process, cellular single-organism, cell part, catalytic activity and binding (Fig. 6). The results of COG functional annotation analysis on the specific and common isoforms in Lm type and normal type castors showed similar to that of GO analysis, and the function of the isoforms remained consistent. COG analysis indicated that the most genes functioned in L (replication, recombination and repair) and R (general function prediction) (Fig. 7).

**Figure 6.**
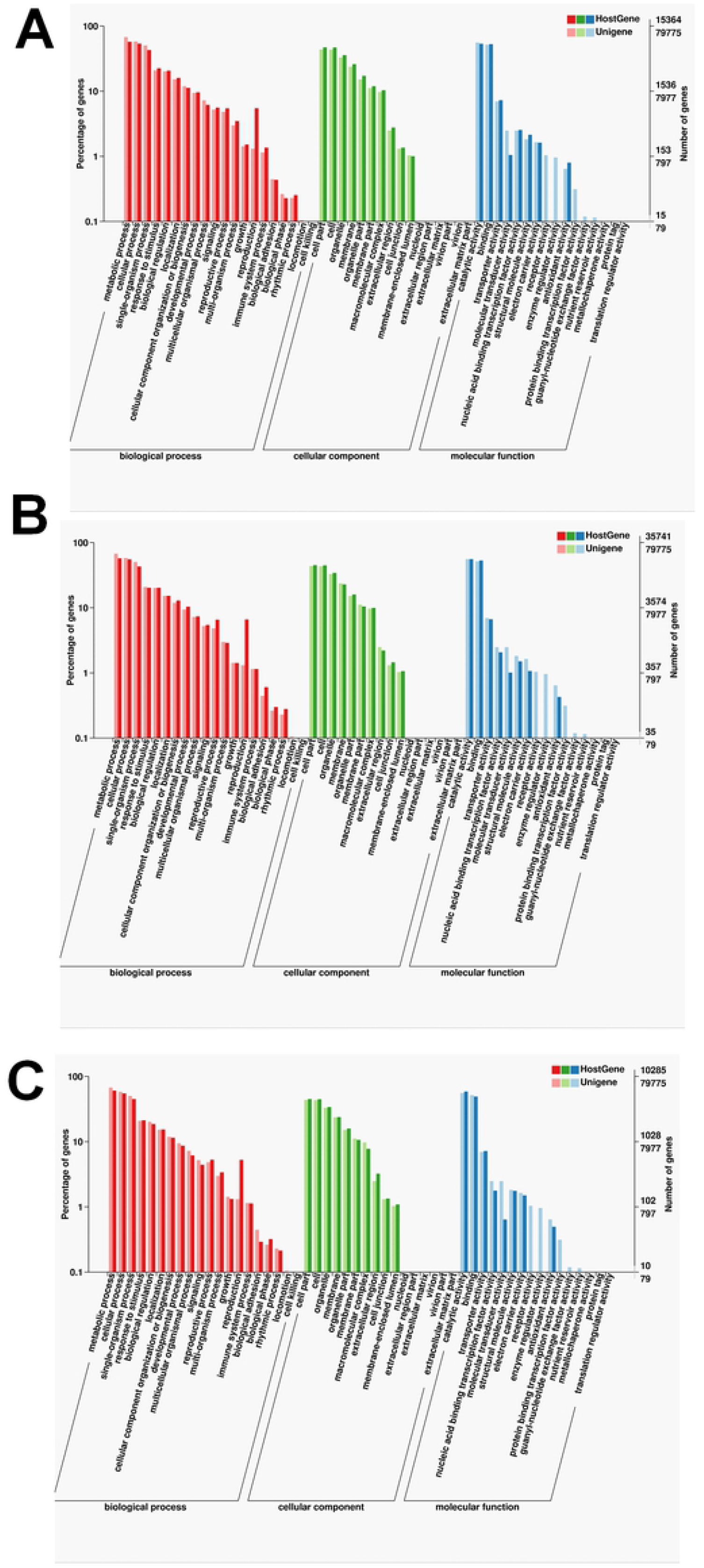
Statistics of the GO classification in two cultivars. **(A)** GO annotation classification statistics common in F01 and F02. (B) GO annotation classification statistics only in F01. (C) GO annotation classification statistics only in F02.

**Figure 7.**
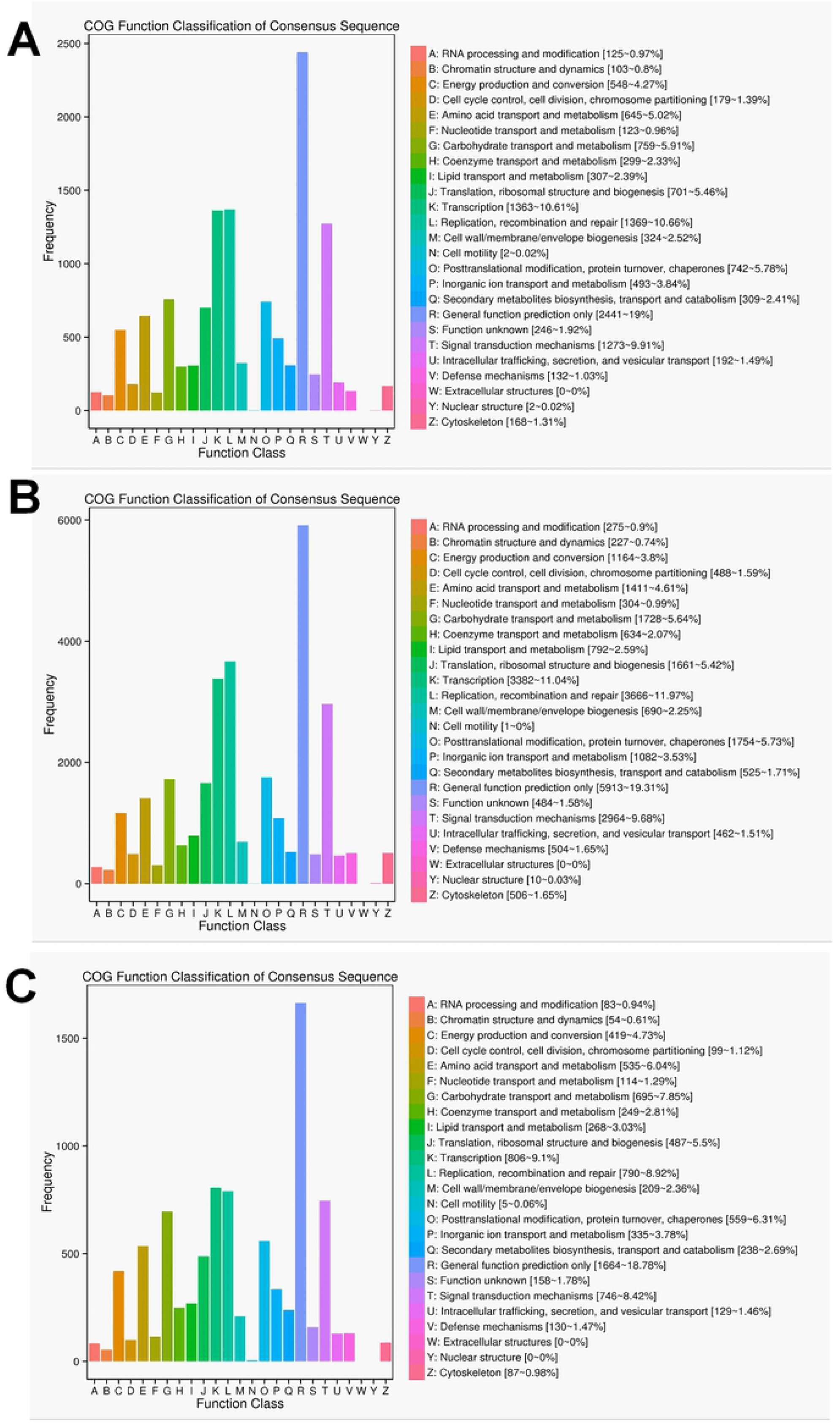
Statistics of the COG classification in two cultivars. **(A)** COG annotation classification statistics common in F01 and F02. (B) COG annotation classification statistics only in F01. (C) COG annotation classification statistics only in F02.

### Comparative analysis of differential gene expression profiling

Full-length sequences were used as a reference genome, and sequences from short-read RNA sequencing were used to conduct differential gene analysis. A total of 2461 genes were found differentially expressed, with 655 up-regulated and 1806 down-regulated. Then these differentially expressed genes (up-regulated genes and down-regulated genes) were classified according to KEGG pathway (Fig. 8). The results showed that up-regulated genes were classified to pathways of ribisome, carbon metabolism, pentose phosphate pathway and biosynthesis of amino acids. Down-regulated genes were involved in plant hormone signal transduction, phenyl propanoid biosynthesis, carbon metabolism, and plant-pathogen interaction.

**Figure 8.**
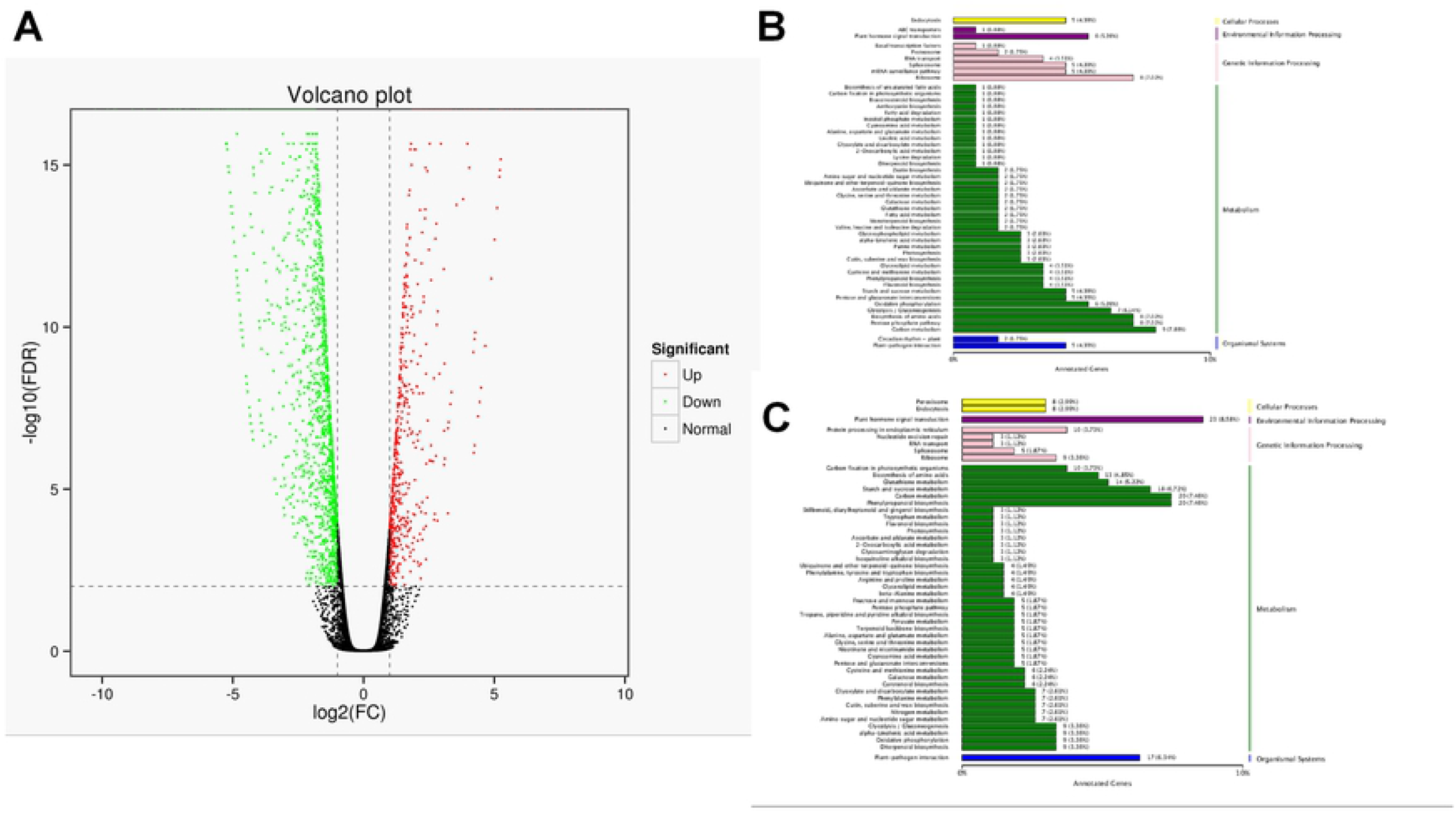
Statistics of DEGs and corresponding function in two cultivars. (A) Volcano plot of DEGs. The green dots represent down-regulated DEGs, the red dots represent up-regulated DEGs, and the black dots represent non-differentially expressed genes. (B) KEGG classification of up-regulated DEGs. (C) KEGG classification of down-regulated DEGs.

## Discussion

### Alternative splicing were involved in phenotypic differences of Lm and normal castor cultivars

In recent years, comparative transcriptome analyses have successfully revealed specific genes responsible for C4 photosynthesis in many grasses, including maize and switch grass [1]. Furthermore, recent studies of the castor transcriptomes are mainly focused on gene expression from short-reads of RNA sequencing[12] [13] which cannot identify alternative gene splice forms [1]. Developing of full-length sequence technology provides a span-new approach to study full-length sequence, alterative splice, gene structure and APA of RNA[14,15]. We thus conducted a comprehensive comparative analysis for two cultivars by using this method. In this work, a total of 76,068 and 44,223 non-redundant transcripts were obtained from the high-quality transcripts of Lm female strain and normal castor, respectively. Among these genes, 51,613 and 20,152 AS events were found in Lm female strain and normal castor, respectively, of which intron retention (IR) was the foremost AS events, with 62.14% and 55.94% in two cultivars, respectively. Its confirmed gene expression and splicing levels might have significant impact on the morphological and other phenotypic differences between two cultivars[16]. Alternative splicing of eukaryotic transcripts is a mechanism that enables cells to generate vast protein diversity from a limited number of genes [17,18]. The mechanism and outcomes of alternative splicing of individual transcripts are well understood [1]. Some studies find that AS regulation is independent or partially independent of transcriptional regulation[19] and implement great function at the early stage of heat response[19,20], useful for future heat sensing and signaling studies [1]. Our new findings of AS provided important information for facilitating castor genome annotation and fully characterization of castor transcriptome.

### Transcription factors and lncRNAs played important role in phenotypic differences of Lm and normal castor cultivars

As the key regulators of transcription, TFs play important role in the physiological regulation of plants [21,22]. Our results (Fig. 3) suggested Rlk-pelle-dlsv, C3H, SNF2 and MYB-related transcription factors were main types in Lm type. Transcription factors of rlk-pelle-dlsv, camk-camkl-chk1, MYB-related bHLH and other types were mainly expressed in normal castor. We speculated that the difference of TFs have a significant effect on the difference in morphology. Similarly, as the important regulator, the number of lncRNA was greatly different: there were 285 lincRNA, 58 antisense-lncRNA, 7 intronic-lncRNA and 166 sense_lncRNA in Lm type, while 60, 22, 3 and 49 in normal castor, respectively. Emerging work has revealed that many lncRNAs regulate gene expression and have great influence on genome stability in plant [23,24]. Studies on Arabidopsis show that lncRNA can serve as molecular sponges and decoys, functioning in regulation of transcription and silencing, particularly in RNA-directed DNA methylation, and in epigenetic regulation of flowering time[25,26]. Many plants reduce expression of some lncRNAs to affect developmental phenotypes or molecular changes [27]. We speculated that these regulators also played important role in growth and development of castor, and contribute significantly to phenotypic differences of Lm and normal cultivars.

### DEGs implement significant function in morphological differences of two cultivars

Using full-length sequences as a reference genome, 2461 differentially expressed genes were found, including 655 up-regulated genes and 1806 down-regulated genes, which was far more than of our previous RNA-seq transcriptome analsysis [1]. In our work, functional analysis showed that proteins encoded by up-regulated genes (655) were classified to ribisome, carbon metabolism, pentose phosphate pathway and biosynthesis of amino acids. Proteins encoded by down-regulated genes (1806) were attributed to plant hormone signal transduction, phenyl propanoid biosynthesis, carbon metabolism, and plant-pathogen interaction. We speculated the differentially expressed genes were the main reason for the difference between the two castors while the specific regulation mechanisms remain unclear.

## Conclusion

This study performed, as our limited knowledge, the first large-scale comparative analysis of the transcriptome by single-molecule long-read sequencing for the Lm type and normal castor cultivars. Comparative analysis of isoforms, transcription factors, lncRNAs and AS in two cultivars were performed systematically. Gene annotation analysis showed that although isoforms were diverse in two cultivars, the implemented functions were similar. Many species-specific differences mainly attributed to small effects at multiple loci probably. However, differences in the expression of genes and alternative splicing events have profound effect on the evolution of major morphological diversification for different individuals in developmental processes. New findings of this study provided invaluable information for facilitating genome annotation and full characterization of transcriptome in two cultivars.

## Declarations

### Ethics approval and consent to participate

This study did not involve any endangered or protected species and followed all relevant ethical guideline. The samples examined in this study were used as agricultural plant in China.

### Consent for publication

Not applicable.

### Availability of data and material

The clean data of RNA-seq were available from National Center for Biotechnology Information Sequence Read Archive database (SRA accession numbers: SRR8662425, SRR8662424).

### Authors’ contributions

ZS designed, supervised the study and wrote the manuscript. WZ performed the study, analyzed data, involved in the writing of the manuscript. YZ involved in data analyses, helped in writing the manuscript. GZ involved in sample collection and preparation, and helped in writing the manuscript. YW and ZH helped perform the analysis, and provided constructive discussions. All authors have read and approved the final manuscript.

## Acknowledgements

The authors appreciate the great help from Professor Guoli Zhu of Tongliao Academy of Agricultural Sciences.

**Supplemental list 1**. KEGG pathway of the genes

**Supplemental Table 1**. Reads of insert (ROI) statistics.

**Supplemental Table 2**. Full-length sequences statistics.

**Supplemental Table 3**. The statistics of different AS events of Lm female strain and normal castor.

## Reference

1. Fan W, Lu J, Pan C, Tan M, Lin Q, Liu W, Li D, Wang L, Hu L, Wang L, Chen C, Wu A, Yu X, Ruan J, Yu J, Hu S, Yan X, Lü S, Cui P (2019) Sequencing of Chinese castor lines reveals genetic signatures of selection and yield-associated loci. Nature communications 10 (1):3418. doi:10.1038/s41467-019-11228-3

2. Tan M, Xue J, Wang L, Huang J, Fu C, Yan X (2015) Transcriptomic Analysis for Different Sex Types of Ricinus communis L. during Development from Apical Buds to Inflorescences by Digital Gene Expression Profiling. Frontiers in plant science 6:1208. doi:10.3389/fpls.2015.01208

3. Podnar J, Deiderick H, Huerta G, Hunicke-Smith S (2014) Next-Generation Sequencing RNA-Seq Library Construction. Current protocols in molecular biology 106:4.21.21-19. doi:10.1002/0471142727.mb0421s106

4. Rogers MF, Thomas J, Reddy AS, Ben-Hur A (2012) SpliceGrapher: detecting patterns of alternative splicing from RNA-Seq data in the context of gene models and EST data. Genome biology 13 (1):R4. doi:10.1186/gb-2012-13-1-r4

5. Buchfink B, Xie C, Huson D (2015) Fast and sensitive protein alignment using DIAMOND. Nature methods 12 (1):59–60. doi:10.1038/nmeth.3176

6. Deng F, Chen SY (2016) dbHT-Trans: An Efficient Tool for Filtering the Protein-Encoding Transcripts Assembled by RNA-Seq According to Search for Homologous Proteins. Journal of computational biology : a journal of computational molecular cell biology 23 (1):1–9. doi:10.1089/cmb.2015.0137

7. Young M, Wakefield M, Smyth G, Oshlack A (2010) Gene ontology analysis for RNA-seq: accounting for selection bias. Genome biology 11 (2):R14. doi:10.1186/gb-2010-11-2-r14

8. Kanehisa M, Furumichi M, Tanabe M, Sato Y, Morishima K (2017) KEGG: new perspectives on genomes, pathways, diseases and drugs. Nucleic acids research 45 (D1):D353–d361. doi:10.1093/nar/gkw1092

9. Leng N, Dawson J, Thomson J, Ruotti V, Rissman A, Smits B, Haag J, Gould M, Stewart R, Kendziorski C (2013) EBSeq: an empirical Bayes hierarchical model for inference in RNA-seq experiments. Bioinformatics (Oxford, England) 29 (8):1035–1043. doi:10.1093/bioinformatics/btt087

10. Jathar S, Kumar V, Srivastava J, Tripathi V (2017) Technological Developments in lncRNA Biology. Advances in experimental medicine and biology 1008:283–323. doi:10.1007/978-981-10-5203-3_10

11. Gong Y, Huang H, Liang Y, Trimarchi T, Aifantis I, Tsirigos A (2017) lncRNA-screen: an interactive platform for computationally screening long non-coding RNAs in large genomics datasets. BMC genomics 18 (1):434. doi:10.1186/s12864-017-3817-0

12. Sood A, Jaiswal V, Chanumolu SK, Malhotra N, Pal T, Chauhan RS (2014) Mining whole genomes and transcriptomes of Jatropha (Jatropha curcas) and Castor bean (Ricinus communis) for NBS-LRR genes and defense response associated transcription factors. Molecular biology reports 41 (11):7683–7695. doi:10.1007/s11033-014-3661-0

13. Sturtevant D, Romsdahl TB, Yu XH (2019) Tissue-specific differences in metabolites and transcripts contribute to the heterogeneity of ricinoleic acid accumulation in Ricinus communis L. (castor) seeds. 15 (1):6. doi:10.1007/s11306-018-1464-3

14. Grabherr MG, Haas BJ, Yassour M, Levin JZ, Thompson DA, Amit I, Adiconis X, Fan L, Raychowdhury R, Zeng Q, Chen Z, Mauceli E, Hacohen N, Gnirke A, Rhind N, di Palma F, Birren BW, Nusbaum C, Lindblad-Toh K, Friedman N, Regev A (2011) Full-length transcriptome assembly from RNA-Seq data without a reference genome. Nature biotechnology 29 (7):644–652. doi:10.1038/nbt.1883

15. Rhoads A, Au KF (2015) PacBio Sequencing and Its Applications. Genomics, proteomics & bioinformatics 13 (5):278–289. doi:10.1016/j.gpb.2015.08.002

16. Wang B, Regulski M, Tseng E, Olson A, Goodwin S, McCombie WR, Ware D (2018) A comparative transcriptional landscape of maize and sorghum obtained by single-molecule sequencing. Genome research 28 (6):921–932. doi:10.1101/gr.227462.117

17. Baralle FE, Giudice J (2017) Alternative splicing as a regulator of development and tissue identity. Nature reviews Molecular cell biology 18 (7):437–451. doi:10.1038/nrm.2017.27

18. Bush SJ, Chen L, Tovar-Corona JM, Urrutia AO (2017) Alternative splicing and the evolution of phenotypic novelty. Philosophical transactions of the Royal Society of London Series B, Biological sciences 372 (1713). doi:10.1098/rstb.2015.0474

19. Siam A, Baker M, Amit L, Regev G, Rabner A, Najar RA, Bentata M, Dahan S, Cohen K, Araten S, Nevo Y, Kay G, Mandel-Gutfreund Y, Salton M (2019) Regulation of alternative splicing by p300-mediated acetylation of splicing factors. RNA (New York, NY) 25 (7):813–824. doi:10.1261/rna.069856.118

20. Keller M, Hu Y, Mesihovic A, Fragkostefanakis S, Schleiff E, Simm S (2017) Alternative splicing in tomato pollen in response to heat stress. DNA research : an international journal for rapid publication of reports on genes and genomes 24 (2):205–217. doi:10.1093/dnares/dsw051

21. Zhou M, Memelink J (2016) Jasmonate-responsive transcription factors regulating plant secondary metabolism. Biotechnology advances 34 (4):441–449. doi:10.1016/j.biotechadv.2016.02.004

22. Olsen AN, Ernst HA, Leggio LL, Skriver K (2005) NAC transcription factors: structurally distinct, functionally diverse. Trends in plant science 10 (2):79–87. doi:10.1016/j.tplants.2004.12.010

23. Wang HV, Chekanova JA (2017) Long Noncoding RNAs in Plants. Advances in experimental medicine and biology 1008:133–154. doi:10.1007/978-981-10-5203-3_5

24. Sun X, Zheng H, Sui N (2018) Regulation mechanism of long non-coding RNA in plant response to stress. Biochemical and biophysical research communications 503 (2):402–407. doi:10.1016/j.bbrc.2018.07.072

25. Zhao X, Li J, Lian B, Gu H, Li Y, Qi Y (2018) Global identification of Arabidopsis lncRNAs reveals the regulation of MAF4 by a natural antisense RNA. Nature communications 9 (1):5056. doi:10.1038/s41467-018-07500-7

26. Liu F, Xu Y, Chang K, Li S, Liu Z, Qi S, Jia J, Zhang M, Crawford NM, Wang Y (2019) The long noncoding RNA T5120 regulates nitrate response and assimilation in Arabidopsis. The New phytologist 224 (1):117–131. doi:10.1111/nph.16038

27. Wang Y, Wang X, Deng W, Fan X, Liu TT, He G, Chen R, Terzaghi W, Zhu D, Deng XW (2014) Genomic features and regulatory roles of intermediate-sized non-coding RNAs in Arabidopsis. Molecular plant 7 (3):514–527. doi:10.1093/mp/sst177

